# Reward impulsivity is linked to addictive-like behaviors towards sweet food and sugary drinks but not fatty food

**DOI:** 10.1101/2025.06.19.660563

**Authors:** Snigdha Mohan, Georgiy Bobashev, Victoria Moiseeva, Anna Shestakova, Ksenia Panidi, Boris Gutkin

## Abstract

Overconsumption of hyperpalatable food is a worldwide problem. Although many previous studies have shown the link between monetary delay discounting and propensity to overconsume food, the studies did not focus on the link between delay discounting and various subcategories of palatable food. In the present study, we investigate the link between the individual propensity to choose smaller immediate rather than larger delayed reward and the degree of problematic eating behavior for five types of palatable food (sweets, sugary drinks, fatty food, salty snacks, and starchy food). The delay discounting task was used to measure propensity to choose immediate rather than delayed rewards, while the degree of addictive-like behavior was measured using the modified Yale food addiction scale (mYFAS) for each food category. Both model-free and model-based analysis showed that there is a significant correlation between the addiction score for sweet food and the tendency to choose smaller immediate rather than larger delayed rewards. A model-based analysis showed the same significant correlation for sugary drinks. However, no such correlation was observed for other food categories (fatty, salty, or starchy food), suggesting that reward impulsivity may play different roles in addictive-like behavior towards various food categories.

## Introduction

Highly palatable food (i.e. food high in salt, fat and sugar) is known to be able to evoke addictive-like behavior ^1,2^. Studies on addictive eating behavior typically focus on energy-dense foods in general, naming high-sugar and/or high-fat products as most common in binge-eating disorder and bulimia nervosa. However, not many studies exist that explore differences in addictive-like behavior towards different food categories containing various combinations of macronutrients, although this issue may have important implications for public policy and clinical interventions. Existing research suggests that both high-fat and high-sugar types of food may be associated with addictive tendencies, but the underlying physiological and psychological mechanisms of this association may differ. In the study of ^3^ among the set of 35 different foods products high in fat content and glycemic load were most likely to be associated with problematic eating behavior. A study with animal models demonstrated that food containing a combination of fat and sugar was most likely to cause activity in the mesolimbic system compared to sugar or fat alone ^4^. Significant differences in demand curves for high-sugar/high-fat versus low-sugar/high-fat products were found using economic methodology ^5^. Individuals were also reported to be willing to pay more for products containing a combination of fat and sugar rather than for fat or sugar alone, controlling for caloric value, and this behavioral response correlated with greater activation of brain reward circuits ^6^. Some studies suggested that sugar and fat may differ in inducing additive-like behavior because of the differences in how they affect dopaminergic system, which leads to withdrawal symptoms for sugary but not fatty food ^7^. Sweet taste was also proposed as a possible mediator of addictive behavior and withdrawal symptoms, although the evidence to support this statement is mixed ^8^.

The aim of the present study is to investigate the relationship between addictive tendencies for five different categories of highly palatable food (sweets, salty snacks, fatty food, sugary drinks, and starchy food) and individual impulsivity measured by a monetary delay discounting task. Delay discounting is a concept that implies the decrease in the value of a reward when the reward is delayed in time. Monetary delay discounting coefficient is currently used in many studies as an indicator of reward impulsivity together with trait impulsivity measured by questionnaires like Barratt Impulsiveness Scale ^9^, and motor impulsivity measured by the performance in a Go/NoGo task ^10^. The choice of smaller and immediate rewards is considered impulsive whereas choosing larger delayed rewards is regarded as a self-controlled choice ^11^. When an individual frequently prefers an immediate smaller reward to a delayed larger reward this indicates a steeper discounting curve and a greater degree of impulsivity ^12^.

Steep delay discounting in the monetary domain has been shown to be associated with various types of addictive behaviors (see ^13^ for a review). The link between delay discounting and addictive behavior towards food has been established as well ^14,15^. A recent review provides evidence that binge-eating disorder is positively associated with reward impulsivity measured by a delay discounting task ^16^. Delay discounting was also suggested to be a behavioral marker of obesity ^17^.

These findings might indicate that the common temporal discounting mechanism stands behind addictive behavior in general. Therefore, if addictive behavior towards sugar stems from the same mechanism we should observe a similar correlation between delay discounting in the monetary domain and addictive behavior towards various foods high in sugar content. In order to separate the effect of fat on sugar-containing food we specifically test for the link between delay discounting and addictive tendencies towards sugary drinks, separately from other food categories, like fatty, starchy, and salty food.

In our study, we asked participants to perform a multi-item monetary choice task to measure their temporal discounting in the monetary domain. Participants also filled in the modified Yale Food Addiction Scale (mYFAS) questionnaire for five food categories. Based on their answers we then analyzed the link between the monetary delay discounting and the addiction score in the five food categories. In the model-free analysis, we investigated the link between the probability to choose immediate rather than delayed reward and the addiction scores for various food categories, without any specific assumptions regarding the delay discounting model. In the model-based analysis, we first use the participants’ responses in the delay discounting task to fit exponential and hyperbolic delay discounting functions, and select the model that fits the behavior best for each particular participant. We then focus on two characteristics of delay discounting: the shape of the discounting curve (exponential or hyperbolic) that best describes each participant’s choices, and their corresponding discounting coefficient. Exponential delay discounting describes consumers whose preferences are time-consistent ^18^. If a consumer prefers an apple over a chocolate bar a week from now, she will still choose an apple over a chocolate bar when asked about what to eat today. However, many people demonstrate hyperbolic discounting, which leads to preference reversals typically observed when the behavior is impulsive. A participant offered an apple versus a chocolate bar in a week from now will make a healthy choice for an apple; if asked to choose what to eat right now they prefer a chocolate bar. Therefore, individuals may be first categorized based on the discounting model (exponential or hyperbolic) that best describes their choices in a delay discounting task. Additionally, individuals with the same discounting type may be further compared based on their discounting coefficient. Those with higher discounting coefficient may be considered more impulsive.

In the model-free analysis, we test the hypothesis that participants with higher addiction scores for sugar-enriched food (such as sweets and sugary drinks) will have higher probability to choose an immediate rather than delayed rewards in a monetary delay discounting task. We found that the hypothesis is confirmed for sweet food, but not for sugary drinks or any other type of food. Participants with higher addiction score for sweets showed higher sensitivity to the amount of money received immediately, with higher immediate amount leading to higher probability to choose immediate option compared to participants with lower addiction scores.

In the model-based analysis, we test two hypotheses. Our first hypothesis holds that individuals who are hyperbolic rather than exponential discounters will be more likely to have higher mYFAS scores for the food categories that contain high levels of sugar. Our second hypothesis suggests that within the group of hyperbolic discounters steeper discounting will be associated with a higher addiction score on the mYFAS scale for sugar-enriched food.

We found that indeed hyperbolic discounters are significantly more likely to have higher addiction scores for sweet food and sugary drinks. Additionally, among hyperbolic discounters higher discounting coefficient (higher impulsivity) was significantly associated with a higher addiction score in the sweet food category. The same tendency was observed for sugary drinks, although it did not reach a conventional statistical significance threshold, possibly because of a lower number of participants showing high addiction scores in this particular category. No statistically significant association was found for fatty, starchy, or salty food.

The present results suggest that reward impulsivity may play a specific role in addictive behaviors towards high-sugar products even when fat is not present in them. The fact that no such relationship was observed for fatty, starchy or salty food may reflect the differences in the addictive mechanisms for sugar-enriched versus other types of food.

## Materials and methods

### Participants

Sixty-seven participants between the age of 18 and 40 years (mean 23y.o., min = 18y.o., max = 38y.o., 52 females) were recruited via a local social media platform and existing university participation database. We excluded participants who self-reported having diabetes, any mental disease, taking prescribed medication, or following any dietary restrictions. The upper age limit was selected to be 40 because taste sensitivity declines with age and is significantly different after the age of 50 ^19^.

The data collection started on 20/08/2020 and ended on 15/09/2020. At a preselection stage, potential candidates filled in an online questionnaire (mYFAS for sweets category). We invited 35 participants who showed no addiction to sweets and 32 who showed at least some addiction to sweets based on this questionnaire. The average age of the participants who are addicted to sweets (PWAS) and non-addicted groups (non-PWAS) was 23.50 and 23.32, respectively.

For all participants, BMI (Body Mass Index) was calculated according to the classification of the World Health Organization ^20^. Body weight and height of the participants were self-reported and used as the measurement to calculate their BMI score. The average BMI score for PWAS and non-PWAS was 22.06 and 21.74, respectively. Our sample comprises individuals who mostly belong within the normal BMI range of 18 to 24.9 with an average score of 22.01. In our sample 70% of the participants had normal BMI, 16% of the participants were underweight, 6% were overweight, and the other 7% were in the obese category.

All procedures were approved by the ethics committee of the HSE university. All experimental procedures were performed in accordance with relevant guidelines and regulations.

### Experimental Protocol

The laboratory experiment consisted of two tasks: completing the mYFAS questionnaire for all five food categories and a computerized temporal discounting task. After completing both tasks participants filled in the BIS/BAS personality scale.

Participants were asked not to eat 2 hours prior to the experiment to keep a similar level of hunger. Because of COVID-19, to maintain safety participants and the experimenter wore masks during the experiment. Before performing the task, all participants signed an informed consent form. Participants were then given written instructions explaining the experimental procedures in detail. Participants were informed that their choices during the delayed discounting task did not have any real-life consequences; however, they were required to make choices as they would in reality. In addition, they were informed that there is no correct or incorrect choice.

Participants then were asked to perform two tasks in a randomized order with instructions written on the screen for each task again. The experimenter was not present to minimize the experimenter’s bias. After completing the experiment, participants were debriefed and paid 250 monetary units (MU) in cash for their participation.

#### (a) Temporal Discounting Task

In our study, we used the multi-item temporal discounting choice task with a staircase procedure^21^. Multi-item choice task has been used in addiction studies for alcohol consumption^22,23^, smoking ^24,25^, and gambling ^26,27^. Similarly, several other studies used the staircase procedure to measure discount rate for a multi-item temporal discounting task ^28,29^.

The procedure was similar to that of ^30^. In each trial, two amounts of hypothetical reward with respect to their time interval would appear on the screen. One reward was immediate and the other was after a delay. The participants had to indicate the option they prefer in every trial by pressing the right or left button. For every trial, a fixation cross was presented for 1 second, after which the choice block between immediate and later rewards was presented. The chosen option was highlighted by a box on the screen for 1 second after a 1.5-second interval. Participants made five choices per block for six time delays (today vs. a delayed day): with 0 days always accounting for present day monetary gain and 2, 14, 30, 60, and 90 days for delayed monetary reward.

The order of each block and time intervals shown on the screen were randomized. Every block started with an option of 200 MU for the immediate option. In all trials, the delayed option was fixed at 400 MU. When the individual chose an immediate reward, the amount of the immediate reward was decreased for the next trial. On the other hand, if the participant chose the delayed reward, the amount of the immediate reward was increased for the next trial. The adjustment amount for the immediate reward decreased with successive choices was chosen following the study by ^31^. In their study, the first adjustment was half of the difference between the immediate and the delayed reward and the next adjustment was half of the previous modification. In our study, the subjects made five choices for each time interval in one block. The immediate reward in each trial represents the subjective value of the delayed reward.

In our experiment for the temporal discounting task we used hypothetical monetary rewards instead of real rewards. The main concern of using hypothetical monetary rewards is that participants might not behave as they would in real life because of lack of motivation. However, studies that have directly compared hypothetical and real monetary rewards did not find any significant difference between the two ^32,33^.

#### (b) mYFAS

Our main instrument in measuring the degree of addiction to sugar enriched food is the mYFAS 2.0 based on seven symptoms of substance dependence using the Diagnostic and Statistical Manual of Mental Disorders Fifth Edition criteria ^34^. mYFAS 2.0 is a 13-item instrument questionnaire that focuses on calculating an addictive score for food based on an individual’s eating pattern of highly palatable foods in the past year. This scale is a shorter version of the YFAS 2.0 questionnaire, which has been proved to have high convergent validity with respect to various eating disorder measures and high discriminant validity with respect to alcohol addiction, behavioral inhibition, and activation measures ^35^. The mYFAS 2.0 questionnaire was shown to have similar properties to YFAS 2.0 in terms of internal consistency and convergent validity having been tested in various populations ^36,37^. Therefore, mYFAS 2.0 is a valid and useful tool to identify individuals with addiction to food and more specifically in our study with sugar dependence.

Participants were asked to fill in the mYFAS 2.0 separately for five food categories: sweets (doughnuts, cookies, ice cream, chocolate, candies), starch (white bread, pasta, rolls, rice), salty snacks (chips, pretzels, crackers), fatty foods (steak, bacon, hamburger, cheeseburger, pizza, french fries), and sugary drinks (soda pop). In the current study, a Russian version of mYFAS 2.0 was created. The validity of the questionnaire translation was tested through back-to-back translation and cognitive interviews.

#### (c) BIS/BAS

The BAS/BIS questionnaire is 24-item self-report measure that identifies two motivational systems: the behavioral inhibition system (BIS), and behavioral activation system (BAS) based on a 4-point Likert scale ^38^. BIS indicates the impulse an individual has to avoid aversive outcomes, whereas BAS is related to the motivation an individual has to approach goal-oriented outcomes. BAS is further divided into three categories—fun seeking, reward responsiveness, and drive. Each category corresponds to measuring the motivation behind finding the novel rewards spontaneously, sensitivity to pleasant reinforcers in the environment, and to follow one’s goals, respectively.

### Data Analysis

#### (1) Model-free analysis

To analyze the behavior in the delay discounting task on a trial-by-trial level we used the logistic regression model with subject-level random effects. We used each participants’ responses on each trial in a delay discounting task as the dependent variable (equal to 1 if delayed option was chosen, and 0 otherwise) to analyze the probability to choose the delayed option. The explanatory variables included the characteristics of the trial (the size of an immediate reward and the time delay to the delayed reward), as well as the addiction scores for each of the five food categories. Additionally, participants’ age, gender, BMI and BIS and Reward Responsiveness scores from the BIS/BAS questionnaire were used as control variables. As the delayed reward amount was kept constant in the task, this variable was not included in the regression model. Importantly, we included the interaction terms of the immediate reward amount and time delay with addiction scores, to test whether the propensity towards addictive behavior is linked to the reward and delay sensitivity in the monetary delay discounting task.

#### (2) Model-based analysis

The model-based data analysis was performed as follows. First, we used 25 choices each participant made in a delay discounting task to fit two models of time discounting—exponential and hyperbolic—with the Bayesian estimation procedure. Then, from these two models for each individual we selected the one that explained their intertemporal choices best (according to the Leave-One-Out information criterion). Therefore, after the estimation procedure, each participant was characterized by two factors: (1) discounting model that fits her choices best (exponential or hyperbolic), (2) estimated discounting coefficient that corresponds to this best-fitting model. We further used regression analysis to test two hypotheses. We hypothesized that participants whose intertemporal choice is best explained by an exponential discounting function will be less likely to demonstrate high addiction scores compared to those with hyperbolic discounting. Second, we expected that for participants in a hyperbolic discounting group higher discounting coefficient (corresponding to steeper discounting) will be associated with higher addiction scores, while no such association will be observed in the exponential discounting group.

Below we describe each step in detail.

##### (a) Discount factor

During the temporal discounting task participants chose between the sooner smaller and the larger later reward on each trial. In accordance with economic theory, we assume that their choice is based on the subjective present value of each option. We considered two possibilities for the delay discounting function—exponential and hyperbolic discounting. In economic theory, if a participant is characterized by the exponential discounting function then their preferences are time-consistent and do not show preference reversals ^39,40^. They are, hence, less likely to demonstrate impulsive behavior such as addiction. By contrast, participants who are characterized by the hyperbolic discounting function are time-inconsistent and are more likely to become addicted to various substances ^41^.

Mathematically, the present subjective value of a reward is equal to the monetary amount multiplied by a discount factor. Therefore, for each participant, we fitted two discounting models, a hyperbolic and an exponential one ^42,43^. Exponential discounting function was assumed to have the following form:

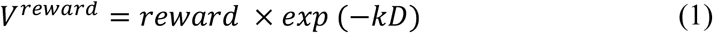

where *D* represents a time delay (in units of time), *k* is a discounting coefficient, and *V^reward^* is the present value of a delayed reward.

The hyperbolic discounting model was a 1-parameter hyperbolic discount function ^44^:

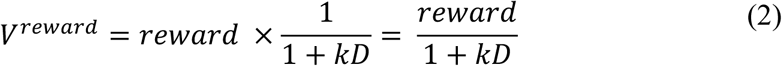

where *D* and *k* represent time delay and a discounting coefficient respectively, and *V^reward^* is the present value of a delayed reward.

We further assumed that a participant’s choice in every trial is determined by the comparison of discounted utilities of each option and some random noise. For the participants’ choice in each trial we assume the random utility model with stochastic errors in *Luce* form ^45^. According to this model, a participant would choose the delayed reward if *log(V_B_) – log(V_A_)>ε*, where *ε* represents a random noise variable, and an immediate reward otherwise. Here, *V_B_* is the present value of a delayed reward, *V_A_* is the present value of an immediate reward, and *ε* represents random error term. We used logs instead of the actual values for better convergence of estimation methods. In accordance with a common modeling practice in economic studies, we assume that random errors in each trial are distributed logistically ^46,47^. Therefore, the probability of choosing the delayed reward is as follows:

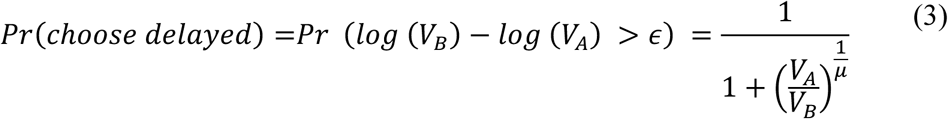

Here, *μ* represents the standard deviation of the noise. If *μ* tends toward zero, then participants would always choose the reward with the highest present subjective value. As *μ* increases the variance of a random error also increases. Hence, the choice between the two options becomes almost random.

To obtain the point estimates of the individual discounting parameter we employed the Bayesian approach. Estimation procedures followed the pipeline described in ^48^. We chose uninformative prior for *k* as a uniform distribution between 0 and 1 and partially informative prior for 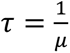 as a uniform distribution between 0 and 10. These parameter constraints are wide enough to cover the values typically found in other delay discounting studies ^32,48^. The obtained posterior distributions did not reach these boundaries, suggesting that the boundaries were not limiting. After estimating the parameters of both exponential and hyperbolic discounting function for each participant, we calculated the Leave-One-Out Cross-Validation information criterion (LOOIC).

For each participant the model with the best fit (lowest LOOIC value) was selected. The type of discounting model that best describes each participant’s preferences was then used as a binary explanatory variable of interest (equal 1 for hyperbolic and 0 for exponential discounting) in the regression analysis of addiction scores.

The estimation procedure was adapted from *hBayesDM* package in R software used to conduct Markov chain Monte Carlo (MCMC) sampling-based inference. The sampling was performed using 4 MCMC chains with 1,000 iterations for warmup and 4,000 iterations in total.

##### (b) Regression analysis

First, we tested the hypothesis that the delay discounting type is linked to addiction to sugar-enriched food. For this purpose, we estimated an ordinal logistic regression model with an addiction score on the mYFAS scale for each food category separately as a dependent variable. Explanatory variables included delay discounting type (exponential or hyperbolic), gender, age, and scores on the BIS and BAS Reward Responsiveness subscales. Because the subscales of the BIS/BAS questionnaire are correlated with each other we used only BIS and Reward Responsiveness scales as independent variables to avoid multicollinearity. These variables were chosen because in many previous studies they demonstrated a significant relation with various kinds of addictions, such as alcohol and marijuana, and gambling behavior ^49–51^.

Second, we tested the hypothesis that for participants with hyperbolic discounting the higher value of the discounting coefficient is associated with increased consumption of sugar-enriched food, while no such relationship is observed for participants best characterized with the exponential discounting model. For this purpose, we estimated the same ordered logit model with discounting coefficient *k* as the main variable of interest. Since the discounting models between participants differ, the values of *k* cannot be pulled together into one variable. Therefore, we included two separate predictor variables. One predictor consists of *k* estimates for the exponential model (*k_exp_*) multiplied by a dummy variable equal to 1 if the best model for a participant is exponential, and zero otherwise (*I_exp_*). The second predictor is constructed in a similar way with *k* values corresponding to the hyperbolic model (*k_hyper_*) multiplied by a dummy variable equal to 1 if the best fitting model for a participant is hyperbolic, and zero otherwise (*I_hyper_*). Since for every participant there is only one best-fitting discounting model (exponential or hyperbolic), the observations for hyperbolic and exponential *k*’s are not included into the regression equation simultaneously. Rather we estimate the effect of *k* on addiction scores separately for each discounting type, while other control variable effects are estimated on the whole sample of participants.

## Results

In our dataset 30 participants showed no tendencies toward addiction to sweets (score 0-1), 20 participants demonstrated medium level of additive tendencies (score 2-5), and 12 participants had severe form of addiction (score 6-13). Of 52 women in our sample, 45% belonged to the non-PWAS category, 35% belonged to the group with moderate addiction to sweets, and 20% had severe level of addiction to sweets. Of 16 men, 75% belonged to the non-PWAS group and both the moderate and severe category of sweet addiction comprised 13% of males each. Discounting model estimates revealed that 47.8% of participants were best characterized with the hyperbolic discounting model and 52.2% with the exponential discounting model. Among the hyperbolic discounters the average estimated discounting coefficient equaled 0.16 (s.e. = 0.033), which corresponds to the devaluation of reward by approximately 14% for the first day of delay. Among the exponential discounters the average estimated *k* equalled 0.093 (s.e. = 0.024), which indicates that for these participants the reward value decreases by 9% per 1 day of delay.

Table 1 presents the estimation results of the regression model in the model-free analysis where no specific assumptions about the shape of the discounting function are made. Here, the dependent variable in each regression model is the probability to choose the larger delayed reward as opposed to the smaller sooner reward in each trial in a delay discounting task. The main explanatory variables of interest are the individual addiction score for various food categories and the interaction term of the addiction score with the immediate reward amount and the time delay.

**Table 1.**
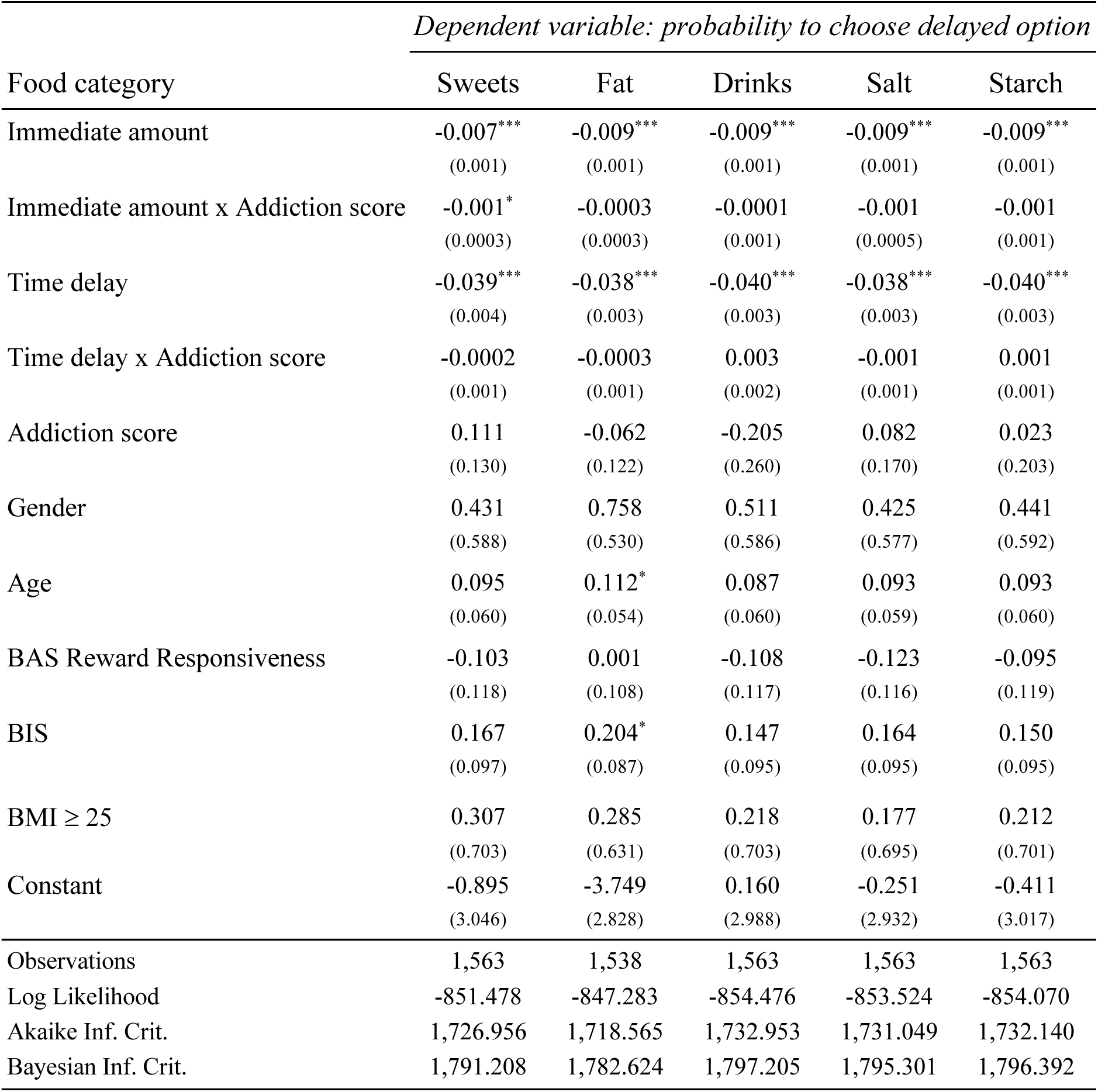
Logistic regression model with subject-level random effects estimation results. Dependent variable in each regression is the probability to choose a delayed reward in each trial in a delay discounting task. Standard errors in parentheses. Stars indicate the following p-value thresholds: *<0.05, **<0.01, ***<0.001.

| Food category | *Dependent variable: probability to choose delayed option* |  |  |  |  |
| --- | --- | --- | --- | --- | --- |
|  | Sweets | Fat | Drinks | Salt | Starch |
| Immediate amount | -0.007***<br>(0.001) | -0.009***<br>(0.001) | -0.009***<br>(0.001) | -0.009***<br>(0.001) | -0.009***<br>(0.001) |
| Immediate amount x Addiction score | -0.001*<br>(0.0003) | -0.0003<br>(0.0003) | -0.0001<br>(0.001) | -0.001<br>(0.0005) | -0.001<br>(0.001) |
| Time delay | -0.039***<br>(0.004) | -0.038***<br>(0.003) | -0.040***<br>(0.003) | -0.038***<br>(0.003) | -0.040***<br>(0.003) |
| Time delay x Addiction score | -0.0002<br>(0.001) | -0.0003<br>(0.001) | 0.003<br>(0.002) | -0.001<br>(0.001) | 0.001<br>(0.001) |
| Addiction score | 0.111<br>(0.130) | -0.062<br>(0.122) | -0.205<br>(0.260) | 0.082<br>(0.170) | 0.023<br>(0.203) |
| Gender | 0.431<br>(0.588) | 0.758<br>(0.530) | 0.511<br>(0.586) | 0.425<br>(0.577) | 0.441<br>(0.592) |
| Age | 0.095<br>(0.060) | 0.112*<br>(0.054) | 0.087<br>(0.060) | 0.093<br>(0.059) | 0.093<br>(0.060) |
| BAS Reward Responsiveness | -0.103<br>(0.118) | 0.001<br>(0.108) | -0.108<br>(0.117) | -0.123<br>(0.116) | -0.095<br>(0.119) |
| BIS | 0.167<br>(0.097) | 0.204*<br>(0.087) | 0.147<br>(0.095) | 0.164<br>(0.095) | 0.150<br>(0.095) |
| BMI ≥ 25 | 0.307<br>(0.703) | 0.285<br>(0.631) | 0.218<br>(0.703) | 0.177<br>(0.695) | 0.212<br>(0.701) |
| Constant | -0.895<br>(3.046) | -3.749<br>(2.828) | 0.160<br>(2.988) | -0.251<br>(2.932) | -0.411<br>(3.017) |
| Observations | 1,563 | 1,538 | 1,563 | 1,563 | 1,563 |
| Log Likelihood | -851.478 | -847.283 | -854.476 | -853.524 | -854.070 |
| Akaike Inf. Crit. | 1,726.956 | 1,718.565 | 1,732.953 | 1,731.049 | 1,732.140 |
| Bayesian Inf. Crit. | 1,791.208 | 1,782.624 | 1,797.205 | 1,795.301 | 1,796.392 |

Estimation results showed that in the delay discounting task participants behaved in an expected way with higher immediate amount and longer time delay leading to lower probability to choose a delayed option in each trial. For each food category, addiction score did not show any significant link with the probability to choose a delayed option, which indicates that high degree of addiction is not associated with an a priori bias towards an immediate or a delayed option. However, the interaction of addiction score with the immediate reward amount was significant for the Sweets category, indicating that participants demonstrating high addiction to sweets had greater sensitivity to the immediate reward amount compared to participants with lower addiction scores to sweets. An increase in the immediate reward amount by 1 monetary unit would lead to greater decrease in the probability to choose the delayed option for participants with higher addiction scores to sweets. No significant association was observed for other food categories.

Table 2 presents the estimation results of regression model in a model-based analysis where participants were separated into exponential and hyperbolic discounters based on the discounting model that best describes their intertemporal choices. Here, the type of discounting was the main explanatory variable of interest.

**Table 2.** Ordered logit model estimation results. Dependent variable in each regression is the individual mYFAS score on the corresponding food category. Standard errors in parentheses. Stars indicate the following p-value thresholds: *<0.05, **<0.01, ***<0.001.

| *Dependent variable:* |  |  |  |  |  |
| --- | --- | --- | --- | --- | --- |
|  | Sweets | Fat | Drinks | Salt | Starch |
|  | (1) | (2) | (3) | (4) | (5) |
| Discounting type |  |  |  |  |  |
| (hyperbolic=1,<br>exponential=0) | 1.144* | 0.868 | 1.245* | 0.390 | 0.649 |
|  | (0.484) | (0.521) | (0.632) | (0.595) | (0.514) |
| Gender | -0.277 | 0.192 | 0.793 | 0.004 | -0.602 |
|  | (0.549) | (0.614) | (0.708) | (0.769) | (0.635) |
| Age | 0.088 | 0.067 | -0.037 | 0.016 | 0.056 |
|  | (0.056) | (0.062) | (0.082) | (0.079) | (0.065) |
| BAS Reward<br>Responsiveness | 0.128 | 0.133 | -0.104 | -0.033 | 0.248 |
|  | (0.118) | (0.130) | (0.143) | (0.140) | (0.135) |
| BIS | 0.217* | 0.039 | 0.088 | 0.270* | 0.103 |
|  | (0.093) | (0.099) | (0.120) | (0.128) | (0.107) |
| BMI ≥ 25 | 0.394 | 0.168 | -0.539 | -0.302 | -1.455 |
|  | (0.585) | (0.647) | (0.881) | (0.903) | (0.900) |
| Observations | 67 | 67 | 67 | 67 | 67 |
| R<sup>2</sup> | 0.254 | 0.072 | 0.110 | 0.109 | 0.181 |
| chi<sup>2</sup> (df = 6) | 19.214** | 4.722 | 6.373 | 6.661 | 12.547 |
| *Note:* |  |  |  | * ** *** p<0.001 |  |

Estimation results suggest that the type of delay discounting is significantly positively associated with an addiction score on the mYFAS scale for *Sweets* and *Drinks* categories. Namely, participants who are hyperbolic rather than exponential delay discounters are more likely to have higher scores on the mYFAS scale. Figure 1 provides further illustration of this result. Additionally, we observe that a higher BIS score is also linked to increased mYFAS scores for *Sweets* and *Salt* food categories.

**Figure 1.**
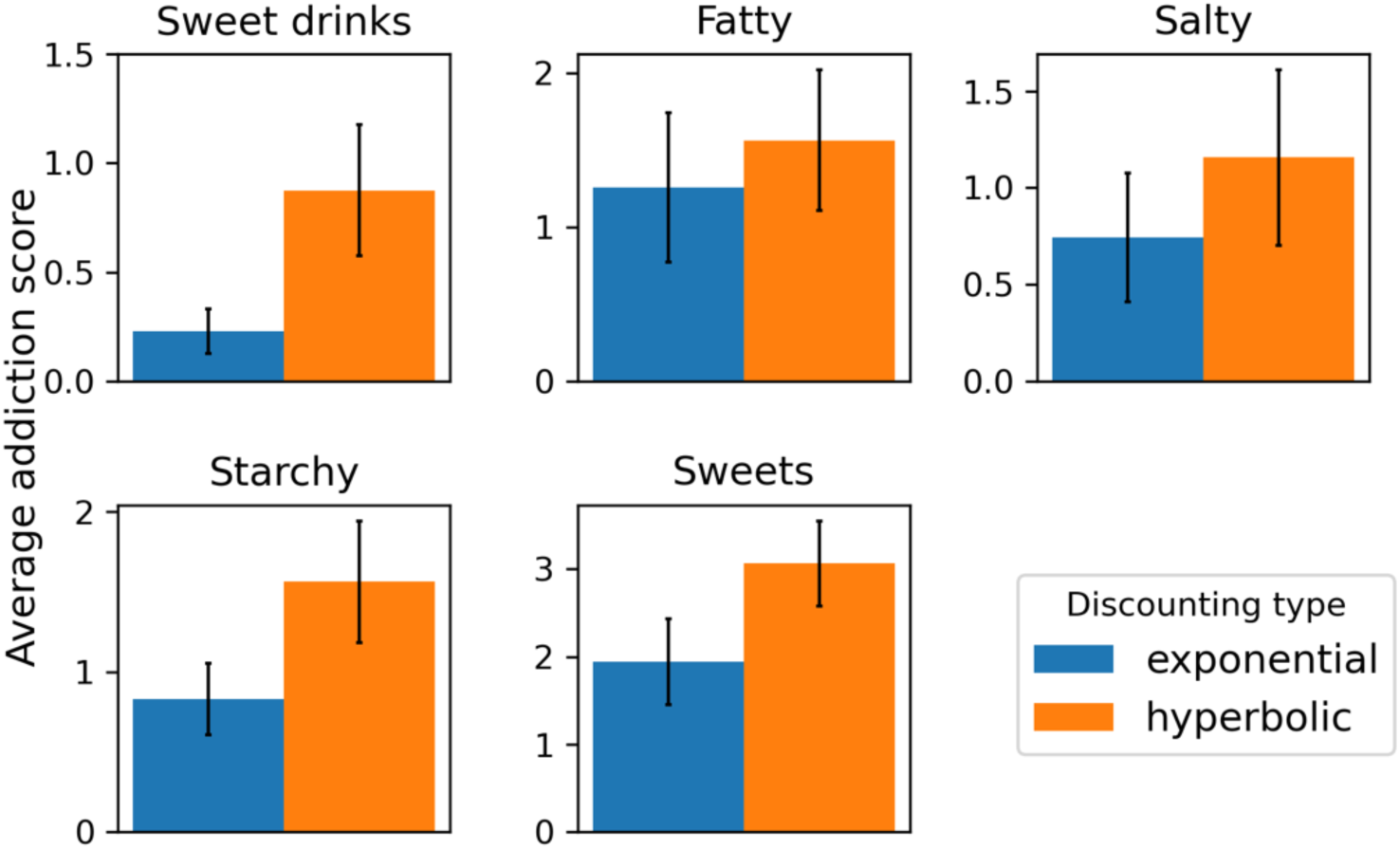
Average addiction score on mYFAS scale in each food category for exponentially and hyperbolically discounting participants. Participants exhibiting hyperbolic discounting demonstrate higher average addiction scores.

Table 3 presents estimation results for the effects of discounting coefficient *k* on addiction scores.

**Table 3.**
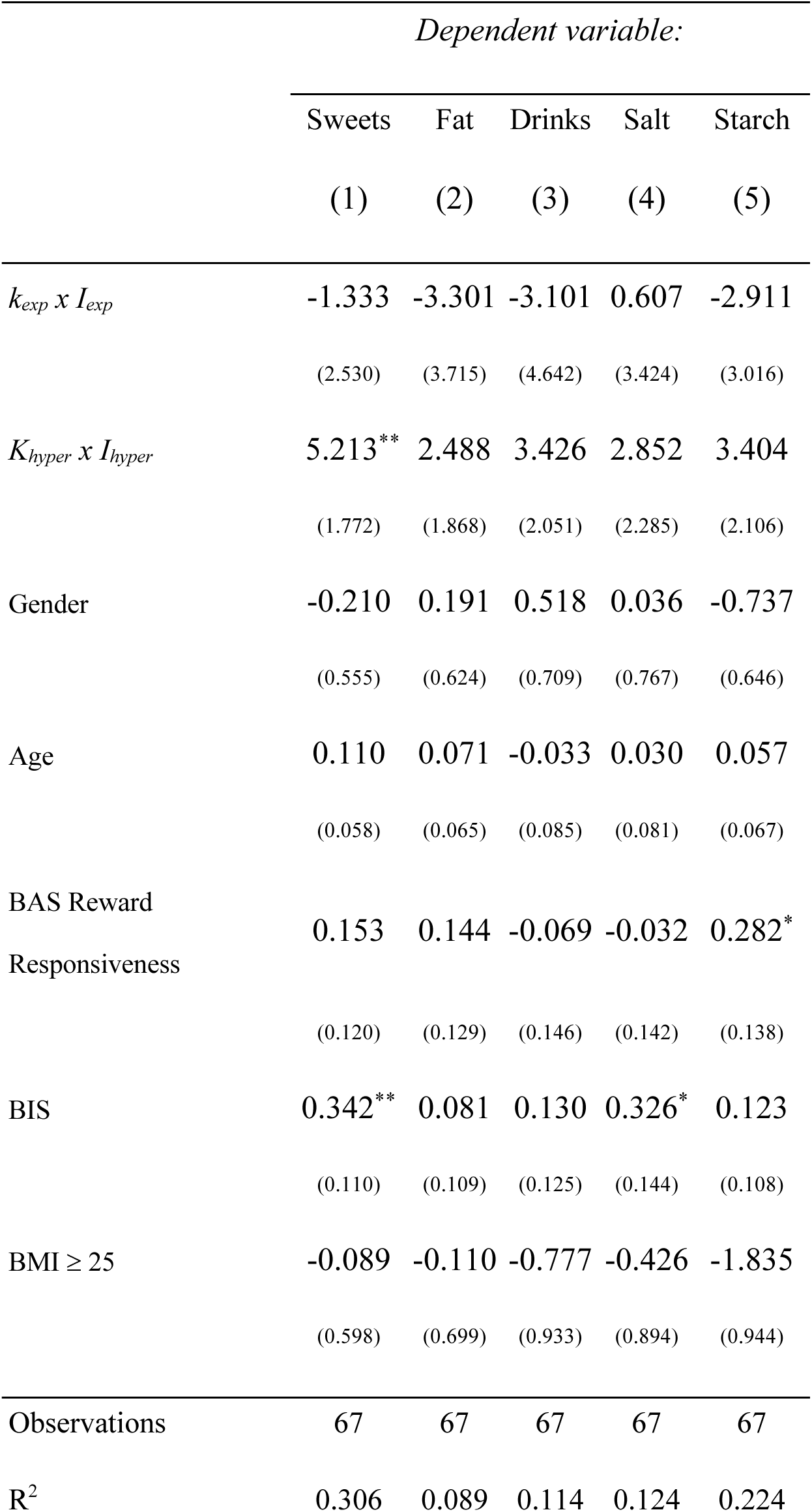
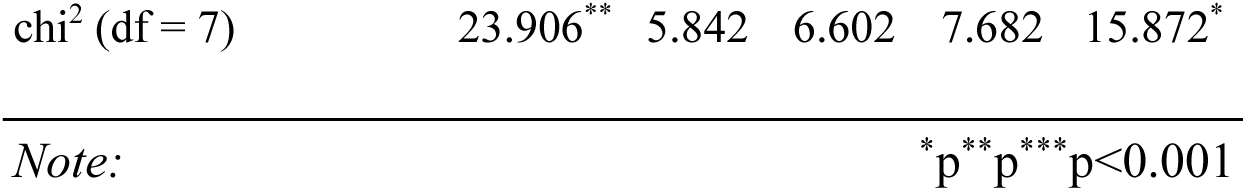
Ordered logit model estimation results. Dependent variable in each regression is the individual mYFAS score on the corresponding food category. *k_exp_* and *k_hyper_* are discounting coefficients estimated assuming exponential and hyperbolic discounting model respectively. *I_exp_* is a dummy variable equal to 1 if the participant’s choices are best fitted with the exponential discounting model, and zero otherwise. *I_hyper_* is a dummy variable equal to 1 if the participant’s choices are best fitted with the hyperbolic discounting model, and zero otherwise. Standard errors in parentheses. Stars indicate the following p-value thresholds: *<0.05, **<0.01, ***<0.001.

As shown in Table 3, for exponential discounters the discounting coefficient is not associated with the addiction score in the *Sweets* category. However, for hyperbolic discounters a larger coefficient *k* (i.e., higher impulsivity in the monetary domain) is significantly linked to a higher addiction score on the mYFAS scale. Interestingly, BIS score is positively associated with the addiction score independently of delay discounting. The same tendency with respect to the delay discounting and mYFAS score is observed in the *Drinks* category; however, it failed to reach significance in our sample (p=0.095). Exponential discounters again do not show any significant correlation with mYFAS scores (p=0.50). No significant association is found in the *Fat*, *Salt,* and *Starch* categories.

## Discussion

In the present study, we explore the association between the behavior in the monetary delay discounting task and addiction to various types of food. Instead of measuring food addiction in general, we use the mYFAS for five different food categories. Both model-free and model-based analysis provides evidence that high degree of addiction to sweet food is associated with more impulsive behavior and a greater sensitivity to monetary rewards in a delay discounting task. In the model-free analysis no assumptions were made on the delay discounting functional form, and participants’ responses were analysed on a trial-by-trial basis. The analysis showed that higher addiction score for sweets led to higher sensitivity to immediate monetary reward and higher probability to make an impulsive monetary choice when this amount is increased. In the model-based analysis, participants who showed hyperbolic instead of exponential discounting were more likely to have high addiction scores for sweets. Additionally, for hyperbolic discounters, higher delay discounting coefficient was also significantly positively associated with the addiction score for sweet food. The same association was found for addiction to sweet drinks in a model-based analysis. However, both, model-free and model-based analysis did not show significant associations between monetary delay discounting and addiction score in other food categories, such as fatty, starchy or salty food.

The fact that the association between delay discounting and addiction score was observed for sugary drinks, which represent products that do not contain fat, suggests that sugar may have high addictive potential independently of other substances like fats. The fact that no such association was observed for fatty food, may suggest that addictive potential of fatty products may be decreased in the absence of sugar, or that addictive mechanisms involved in addictive-like behaviors towards fatty food are distinct from those involved in addictive behavior towards sugar-enriched food.

## Data availability

Complete dataset used in this study is freely available in the repository: https://github.com/openaccessdata/sugar_addiction

## Funding

This article is an output of a research project implemented as part of the Basic Research Program at the National Research University Higher School of Economics (HSE University).

## Author contributions

SM: conceptualization, data curation, formal analysis, drafting original writing;

GB: supervision, reviewing and editing;

VM: project administration, funding acquisition;

AS: supervision, project administration, funding acquisition;

KP: conceptualization, supervision, formal analysis, reviewing and editing;

BG: supervision, reviewing and editing.

## Notes

https://github.com/openaccessdata/sugar_addiction

### Competing Interest Statement

The authors have declared no competing interest.

### Summary of Updates

Funding information added in the end of the manuscript.

